# The role of single occupancy effects on integrase dynamics in a cell-free system

**DOI:** 10.1101/059675

**Authors:** Georgios Artavanis, Victoria Hsiao, Clarmyra A. Hayes, Richard M. Murray

**Affiliations:** Division of Biology and Biological Engineering, California Institute of Technology, Pasadena, California, USA

## Abstract

Phage integrase-based circuits are an alternative approach to relying on transcriptional and translational repression for biomolecular circuits. Previous research has shown that circuits based on integrases can perform a variety of functions, including counters, Boolean logic operators, memory modules and temporal event detectors. It is therefore essential to develop a principled theoretical and experimental framework for the design, implementation and study of such circuits. One of the fundamental questions that such a framework should address concerns the functionality limitations and temporal dynamics of the integrases as regulatory elements. Here, we test the functionality of several large serine integrases from a recently published library in a cell-free transcription-translation (TX-TL) platform. Additionally, we use a combination of experimental data and models to investigate integrase dynamics as a function of enzyme concentration and number of binding sites.

We report that sequestration of integrase molecules, either in the form of monomers or dimers, by the integrase's own binding sites dominates integrase dynamics, and that the delay in the activation of the reporter is negatively correlated with integrase plasmid concentration. We have validated our sequestration hypothesis by building a model with MATLAB’s SimBiology toolbox, and running simulations with various integrase and binding sites concentrations. The simulation results qualitatively match the experimental results, and offer further insights into the system.

## Introduction

One of the main goals of synthetic biology is to unveil the design principles of biomolecular circuits, as well as to use this knowledge to engineer circuits with desirable properties. So far, the main approach has been to rely on transcriptional and translational regulation in order to manipulate the characteristics and behavior of biomolecular circuits.

A novel, alternative paradigm is emerging, in which phage integrases are used as the main regulatory elements of biomolecular circuits. Integrases are proteins that modify DNA and are often used by viruses as part of the lysogenic machinery. Previous research has shown that circuits based on integrases combined in novel ways (possibly with nested recognition sites) can perform a variety of functions, including counters [1], Boolean logic operators [2, 7], memory modules [3] and temporal event detectors [4]. In order to take full advantage of the range of possible functions that can be achieved by integrase-based circuits, it is necessary to develop a principled theoretical and experimental framework for the design, study and implementation of these circuits. Such a framework should address functionality limitations, temporal dynamics, design principles and experimental protocols for the implementation of phage integrases as regulatory elements of engineered circuits.

One of the fundamental questions that such a framework should address concerns the functionality limitations and temporal dynamics of large serine integrases as regulatory elements. We have tested the functionality of several large serine type integrases from a recently published library [5] in a well-mixed cell-free transcription-translation (TX-TL) platform. In addition, we have explored experimentally and through mathematical modelling and simulations how integrase dynamics depends on the concentration of integrase and that of its binding sites.

The TX-TL platform is a cell-free expression system consisting of bacterial cell extract (lysate) and buffer; It allows implementation of biomolecular circuits quickly and efficiently as plasmid systems or even as pieces of linear DNA [6]. We measure the outputs of the implemented circuits through time-course bulk fluorescence readings.

To study the functionality and dynamics of each integrase, we used a two plasmid system developed in [5]. The first plasmid encodes an integrase whose expression is driven under an arabinose-inducible promoter. The second plasmid encodes the GFP reporter, under the control of a constitutive promoter, that is flanked by the integrase’s recognition sites. The reporter is initially not expressed, as the GFP coding region is inverted. Once the integrase has been produced, it binds to its recognition sites flanking the reporter and flips the reporter (GFP), allowing its expression. A phage integrase recognizes two attachment sites, one from the phage (attP) and one from the host (attB). For DNA inversion, two integrase monomers bind to each attachment site, followed by DNA looping and recombination of the sites. The new sites, attL and attR, each comprise half the attP and half the attB sequence (Figure 1). The reverse reaction only happens in the presence of a cognate excisionase, otherwise DNA inversion is strictly unidirectional [8].

In this report, we show that sequestration of integrase molecules, either in the form of monomers or dimers, by its own binding sites dominates integrase dynamics, and that the delay in the activation of the reporter is negatively correlated with integrase plasmid concentration. We validate this sequestration hypothesis by building a model with MATLAB’s SimBiology toolbox, and running simulations with various integrase and binding site concentrations. The simulation results qualitatively match the experimental results, and offer further insights into the system. By demonstrating that large serine type integrases are functional in a cell-free system and describing the underlying dynamics, our work paves the way for rapid prototyping, testing and debugging of integrase-based biomolecular circuits.

**Figure 1:**
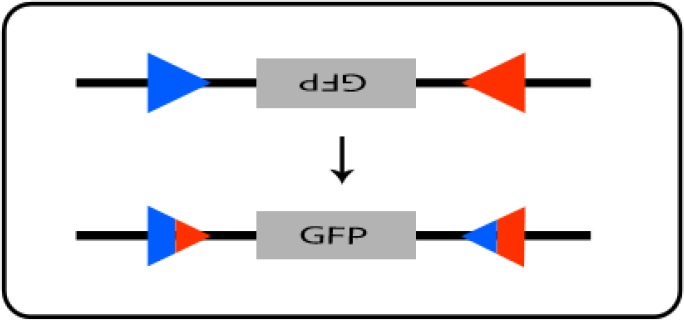
*Schematic of the integrase recombination action. Figure based on Yang L. et al. (2014)*

## Results and Discussion

In [5], a library of 13 previously uncharacterized serine integrases were characterized and shown to be orthogonal *in vivo*. We tested this library, and report that several of these integrases are functional in the TX-TL platform (Int8, Int10, Int12, Int13). Because of energy limitations of the TX-TL platform, we chose to concentrate on the integrase with the least activation delay of the group, Int8, to explore the range of dynamic responses possible. We include preliminary data from the testing of several other integrases at the end of this report.

In Figure 2, we present the output of the system for different integrase (Int8) and reporter plasmid input concentrations, measured by time-course bulk fluorescence measurements. Each experimental trace contains data from 3 replicates. We found that delays in the activation of the reporter are negatively correlated with the concentration of integrase (Fig. 3 A, B, C). We also observed that the concentration of the reporter does not significantly alter the dynamics in the cases of the two highest integrase plasmid concentrations (Fig. 3 B, C), and we hypothesize that this is because of binding site saturation with integrase monomers or dimers. In order to study dynamics out of this saturation regime, it will be necessary to overcome energy limitations of the TX-TL platform [9]. Overall, the steady state fluorescence of the reporter is lower with lower integrase plasmid concentration, since lower concentration of integrase causes a bigger delay in the reporter activation and hence greater energy depletion over time.

The data suggest a single occupancy effect dominating the initial dynamics: in our well-mixed system, after integrase molecules are produced, they bind reporter plasmids randomly, either as monomers or dimers. The probability that a fixed reporter plasmid has 4 monomers bound to it, the number required for DNA recombination, is small initially and gets larger as more integrase monomers are produced. Thus, reporter plasmids are not flipped sequentially, but rather simultaneously, resulting in the sharp initial activation slopes. The effect is enhanced by the release of integrase molecules after site recombination occurs.

**Figure 2:**
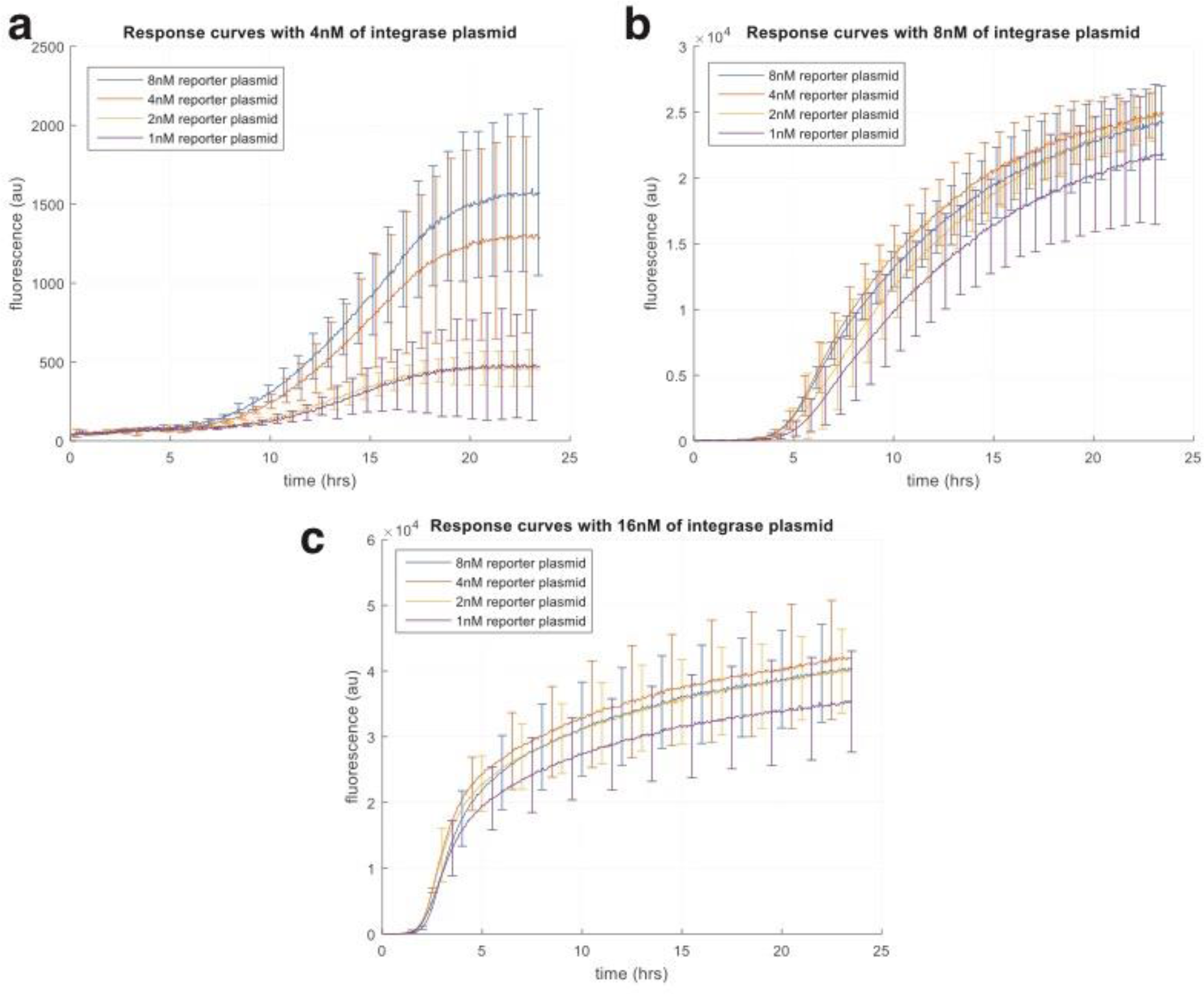
*TX-TL experimental data with varying int8 integrase plasmid concentration and int8 reporter concentrations. (a) Response curves with 4nM of integrase plasmid. (b) Response curves with 8nM of integrase plasmid. (c) Response curves with 16nM of integrase plasmid*.

We validated our sequestration hypothesis by building a model with MATLAB’s SimBiology toolbox, and running simulations with various integrase and binding site concentrations. The simulation results qualitatively match the experimental results, and offer further insights into the system. Interestingly, our model predicts that dimerization rates of integrase molecules prior to site-specific binding do not significantly affect the overall dynamics. We include the SimBiology model file in the supplementary data.

In Figure 3, we present qualitative simulations of the model with varying integrase and reporter plasmid concentrations. We simulated integrase plasmid concentrations of 1nM, 10nM, and 100nM (Fig. 3 A, B, C, respectively).

**Figure 3:**
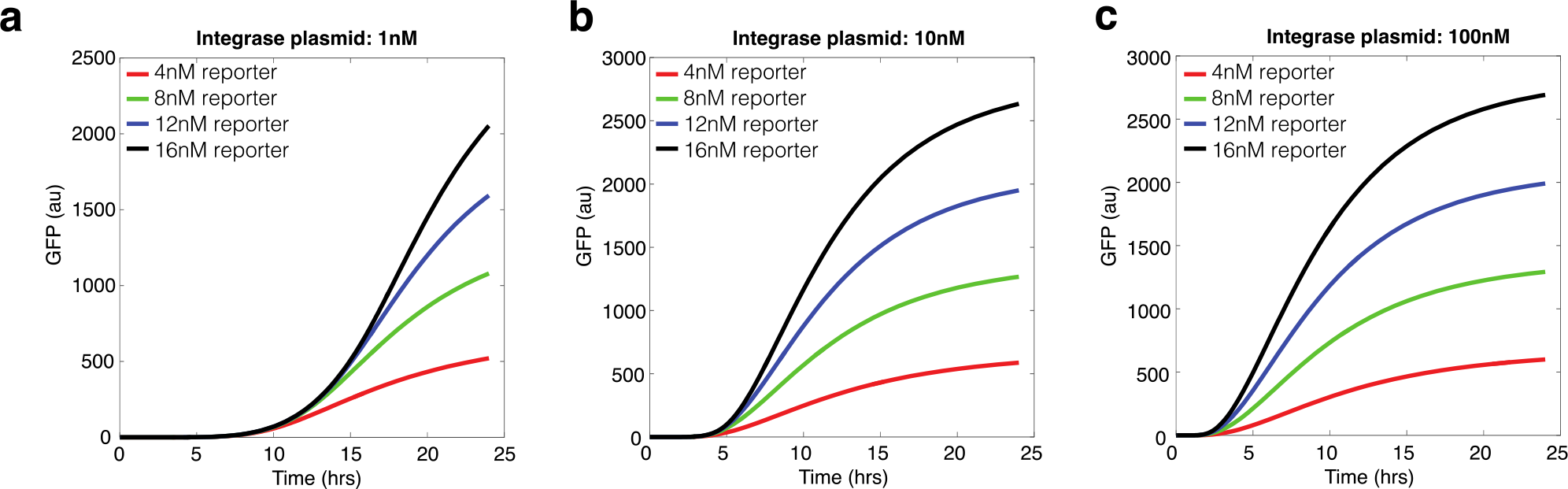
*Simulation results for varying integrase and binding site concentrations. a) 1nM of integrase plasmid. b) 10nM of integrase plasmid. c) 100nM of integrase plasmid. The simulations show that the delay in the activation of the reporter is negatively correlated with the concentration of integrase*.

In our model, plasmid concentrations were used as multipliers of integrase transcription and translation. After integrase molecules are produced, they form dimers that bind the integrase’s target sites. The initial concentration of non-recombined binding sites is the modeled concentration of the reporter plasmid. When a reporter plasmid has an integrase dimer bound to both target sites, an intermediate complex is formed between the DNA and the two integrase dimers, that represents the looping stage of recombination. Finally, the DNA is flipped and the two integrase dimers are released, allowing the reporter’s transcription and translation. All chemical reactions follow the law of mass action.

In both the experimental data and the simulations, at a constant integrase concentration, higher reporter plasmid concentrations result in smaller delays in the recombination and activation of the reporter, which cannot be explained in terms of the single occupancy dynamics alone: According to the single occupancy dynamics, higher reporter plasmid concentrations (and hence higher binding site concentrations) would slow the dynamics, by *‘*diluting out*’* the number of available integrase molecules, which is fixed. However, the higher binding site concentrations also increase the rate of the reactions taking place, such as the binding of integrase molecules on them. In Figure 4, higher concentrations of binding sites result in earlier and higher binding of the integrase dimers and hence in earlier flipping of the GFP reporter. Thus, the single occupancy effects demonstrate themselves with varying concentrations of integrase, but at a fixed concentration of integrase, varying the concentration of the reporter also creates other effects that oppose the single occupancy effects.

**Figure 4:**
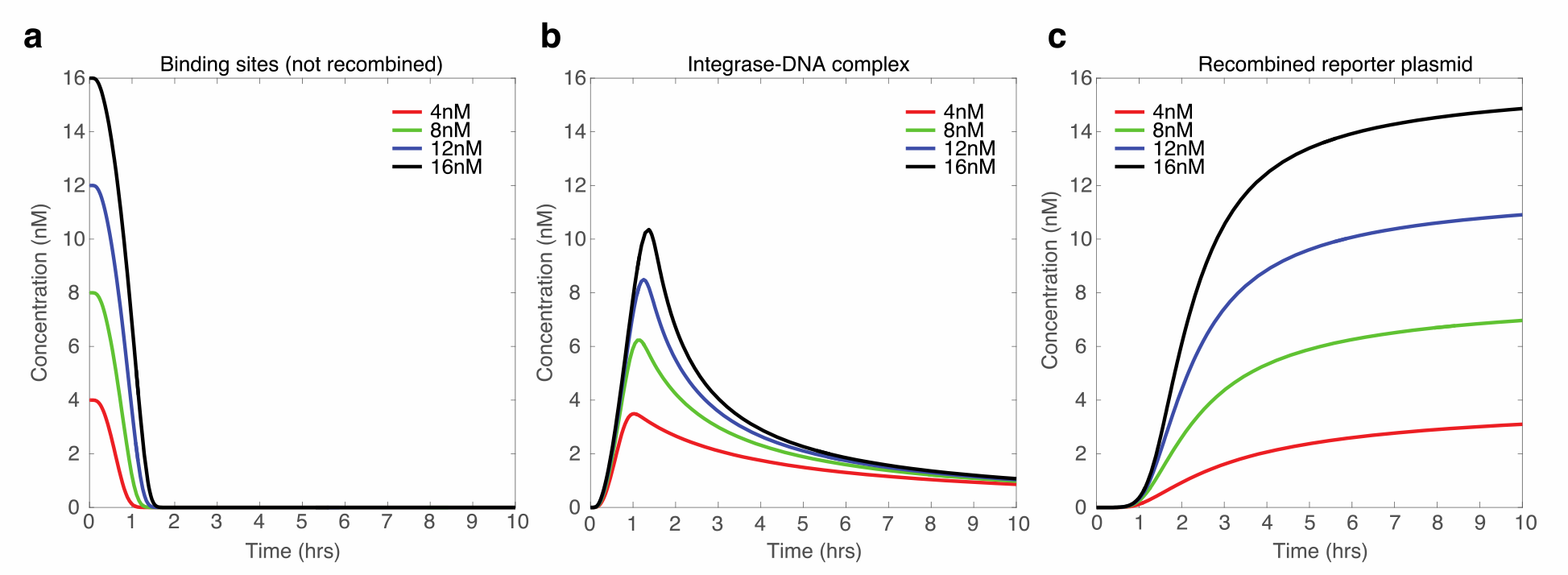
*Higher concentrations of binding sites increase the rate of reactions taking place. Thus, higher concentrations of reporter result in earlier and higher binding of the integrase dimers to the binding sites and hence in earlier activation of the reporter*.

## Conclusions and Future Work

We have demonstrated functionality of several large serine integrases in a cell-free environment, and explored the dynamics of their recombination action. Our results suggest that single occupancy effects of the integrase attachment sites dominate the initial dynamics, resulting in delays in the reporter activation when integrase concentration is lower. A mathematical model of the system has confirmed our hypothesis, and allowed further insights in the system’s dynamics. Thus, we can take advantage of the efficiency and scalability of the TX-TL platform to rapidly test and debug integrase-based biomolecular circuits.

To develop a comprehensive theory of integrase-based circuits, it will be important to investigate the dynamics of the integrases further, and how it depends on environmental factors (e.g. temperature). To study dynamic equilibrium steady states, such as bidirectional flipping in the presence of the cognate excisionase, energy limitations of the TX-TL platform must be overcome, perhaps with the use of microfluidic systems replenishing energy at a constant rate. Finally, a detailed model of the energy-dependency of the several intermediate steps from integrase production to DNA inversion would be useful in determining how costly are integrase-based circuits, which should be taken into consideration when implementing them *in vivo*.

**Figure 5:**
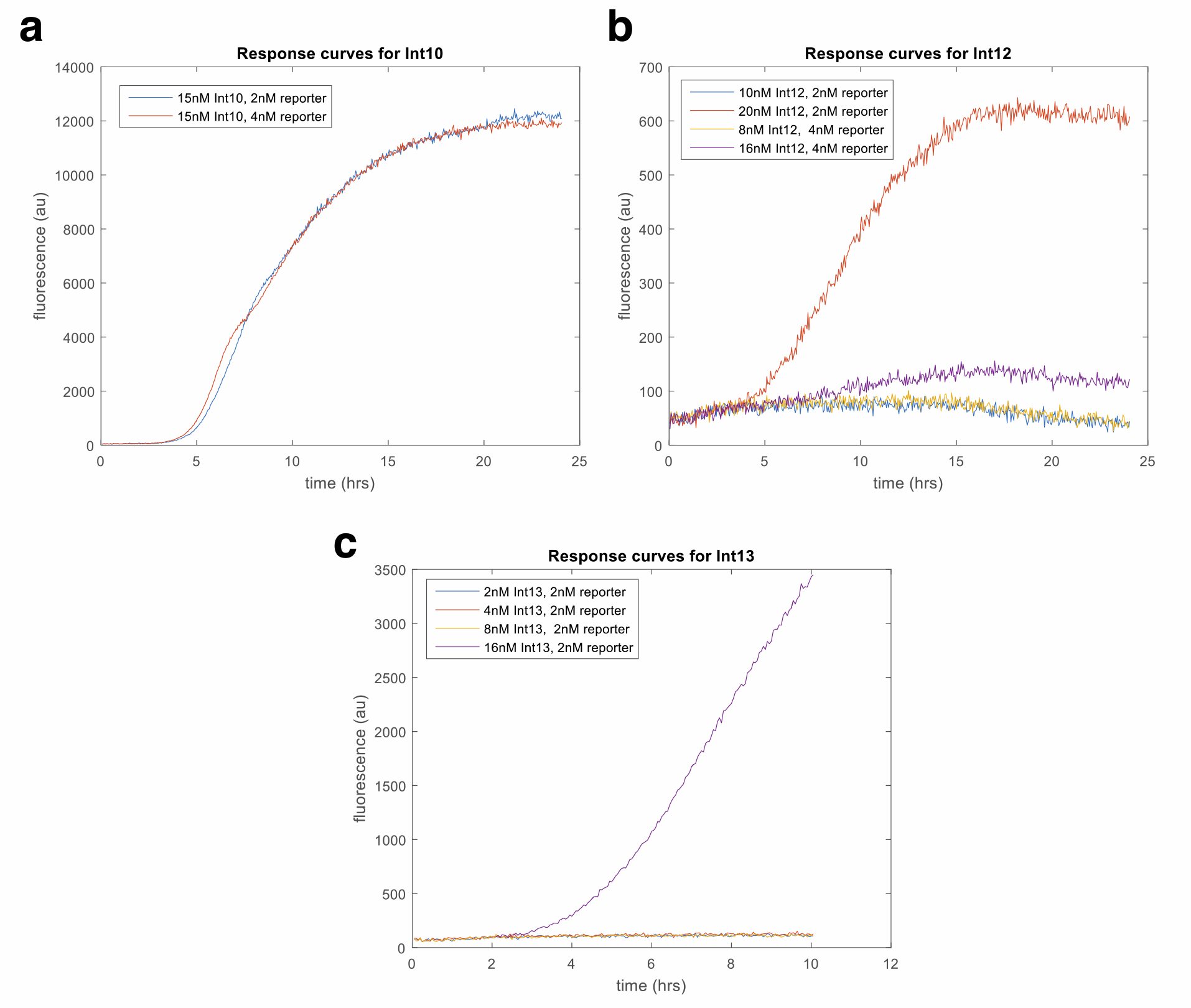
Response curves for Int10, Int12, Int13, illustrating functionality of these integrases in the cell-free TX-TL platform. Each integrase has its own characteristic activation delay and dynamical range.

## Materials and Methods

### Plasmids

The plasmids used were obtained from the Addgene repository (Addgene IDs 60563-60584).

### TX-TL Experiments

Cell extract and buffer were prepared following the protocol described in [6]

### Plate Reader Measurements

Plate reader data were collected on a Biotek H1MF machine using the kinetic read feature, at 29C with no shaking. RFP measurements were taken at excitation/emission of 580/610 with gain 140, GFP measurements were taken at excitation/emission of 485/528 with gain 75.

### Model Implementation

The model was implemented using the SimBiology toolbox in MATLAB and using the ode15s solver (see supplementary file for the model).

## Acknowledgements

G.A. is supported by Caltech BBE Divisional Funding. V.H. is supported by the U.S. Department of Defense (DoD) through the National Defense Science & Engineering Graduate Fellowship (NDSEG) Program. Research supported in part by the Institute for Collaborative Biotechnologies through grant W911NF-09-0001 from the U.S. Army Research Office. The content of the information does not necessarily reflect the position or the policy of the Government, and no official endorsement should be inferred.

## References

[1] A. E. Friedland, T. K. Lu, X. Wang, D. Shi, G. Church, and J. J. Collins. Synthetic gene networks that count. Science, 324(5931):1199–1202, 2009.

[2] P. Siuti, J. Yazbek, and T. K. Lu. Synthetic circuits integrating logic and memory in living cells. Nature Biotechnology, 31:448–452, 2013.

[3] J. Bonnet, P. Subsoontorn, and D. Endy. Rewritable digital data storage in live cells via engineered control of recombination directionality. PNAS, 109:8884–8889, 2012.

[4] V. Hsiao, Y. Hori, P. W. K Rothemund, and R. M. Murray. A population-based temporal logic circuit for timing and recording chemical events. Molecular Systems Biology, 12(5), 869, 2016.

[5] L. Yang, A. A. K Nielsen, J. Fernandez-Rodriguez, C. J. McClune, M. T. Laub, T. K. Lu, and C. A. Voigt. Permanent genetic memory with >1-byte capacity. Nature Methods, 2014.

[6] Sun Z.Z., Hayes C.A., Shin J., Caschera F., Murray R.M., Noireaux V. Protocols for implementing an Escherichia coli based TX-TL cell-free expression system for synthetic biology. J. Vis. Exp. 2013: e50762.

[7] J. Bonnet, P. Yin, M. E. Ortiz, P. Subsoontorn, and D. Endy. Amplifying genetic logic gates. Science, 340(6132):599–603, 2013.

[8] Groth AC, Calos MP. Phage integrases: biology and applications. J. Mol. Biol. 335:667–678, 2004.

[9] Noireaux V, Bar-Ziv R, Libchaber A. Principles of cell-free genetic circuit assembly. PNAS. 100:12672–12677, 2003.

